# Systematic analysis of YFP gene traps reveals common discordance between mRNA and protein across the nervous system

**DOI:** 10.1101/2022.03.21.485142

**Authors:** Joshua S Titlow, Maria Kiourlappou, Ana Palanca, Jeffrey Y Lee, Dalia S Gala, Darragh Ennis, Joyce J S Yu, Florence L Young, David Miguel Susano Pinto, Sam Garforth, Helena S Francis, Finn Strivens, Hugh Mulvey, Alex Dallman-Porter, Staci Thornton, Diana Arman, Aino I Järvelin, Mary Kay Thompson, Ilias Kounatidis, Richard M Parton, Stephen Taylor, Ilan Davis

**Affiliations:** Department of Biochemistry, University of Oxford, South Parks Road, Oxford OX1 3QU; Wellcome Leap - Delta Tissue Program; Departamento de Anatomía y Biología Celular, Universidad de Cantabria, Cantabria; Weatherall Institute for Molecular Medicine, University of Oxford, Oxford

## Abstract

While post-transcriptional control is thought to be required at the periphery of neurons and glia, its extent is unclear. Here, we investigate systematically the spatial distribution and expression of mRNA at single molecule sensitivity and their corresponding proteins of 200 YFP trap protein trap lines across the intact *Drosophila* nervous system. 98% of the genes studied showed discordance between the distribution of mRNA and the proteins they encode in at least one region of the nervous system. These data suggest that post-transcriptional regulation is very common, helping to explain the complexity of the nervous system. We also discovered that 68.5% of these genes have transcripts present at the periphery of neurons, with 9.5% at the glial periphery. Peripheral transcripts include many potential new regulators of neurons, glia and their interactions. Our approach is applicable to most genes and tissues and includes powerful novel data annotation and visualisation tools for post-transcriptional regulation.

**Brief outline:** A novel high resolution and sensitive approach to systematically co-visualise the distribution of mRNAs and proteins in the intact nervous system reveals that post-transcriptional regulation of gene expression is very common. The rich data landscape is provided as a browsable resource ([link](https://zegami.com/collections/public-61f9786b8e9607c337c5f46c?list.isThisgenescreened.any=Yes&fc=isThisgenescreened&pan=SNAPSHOT_PANEL&view=grid)), using Zegami, a cloud-based data exploration platform ([link](https://zenodo.org/record/6379926/files/Zegami_manual_Zenodo_20220323S.pdf?download=1)). Our solution provides a paradigm for the characterisation of post-transcriptional regulation of most genes and model systems.

**Highlights:**

- 196/200 (98%) *Drosophila* genes show discordant RNA and protein expression in at least one nervous system region
- 137/200 (68.5%) mRNAs are present in at least one synaptic compartment
- Novel localised mRNA and protein discovered in periphery of glial processes
- New paradigm for analysis of post-transcriptional regulation and data exploration

## Introduction

Neurons are the most extremely polarised cell type in multicellular organisms, with many distinct peripheral sites that have to act independently, namely dendritic and axonal synapses. It has been generally accepted that delivering molecules to the periphery of cells involves mRNA localisation in fibroblast cells (Sundell and Singer 1990). But in neurons, the periphery is a large distance from the cell body, requiring long distance transport and localised translation of peripheral transcripts in order to regulate protein levels at the synapses (Holt, Martin, and Schuman 2019). Conclusive examples of mRNA transport and localised translation in dendrites and oligodendrocytes have been known for decades (Carson, Kwon, and Barbarese 1998; Steward et al. 1998). However, axonal localisation and local translation has been easier to discover in developing axons and slower to be elucidated in mature axons. Nevertheless, convincing examples have been known for some time (Jung, Yoon, and Holt 2012), despite mRNA being only found at low concentrations at or near the distant synapses of axons. Efforts to address the relative proportion of mRNAs that are locally translated at synapses has led to some outstanding studies, showing that localised mRNA and local translation are common (Hafner et al. 2019). Such data have been complemented with specific conclusive experiments in intact nervous systems (D. O. Wang et al. 2009). However, it is not known what the relative contribution of local translation versus nuclear transcription and protein transport are in the diverse cell types of an intact functional mature nervous system.

A hallmark of post-transcriptional regulation is that the distribution of individual species of protein and mRNA are discordant, or uncorrelated. Such discordance is most obviously manifested in a lack of correlation between the levels of mRNA expression and protein levels across distinct cell types in a tissue, through mRNA stability differences or variations in the rates of translation. However, post-transcriptional regulation can also manifest itself within a cell, so that a protein is localised to a distinct site from the mRNA that encodes it. Many mechanisms can lead to intracellular protein and mRNA discordance, including localised translation, mRNA degradation or intracellular transport of protein or mRNA (Mofatteh and Bullock 2017). To date, systematic characterisation of discordance between protein and mRNA have not been carried out across a whole intact nervous system or any other complex tissue. The advent of single cell transcriptomics (Aldridge and Teichmann 2020) and spatial transcriptomics (Marx 2021) has been a major transformational step. However, although single cell proteomics and high-coverage imaging mass spectrometry are on the horizon (Marx 2019), the methods currently lack sufficient sensitivity or coverage, have limited resolution, and are unable to multiplex RNA and protein detection at substantial scale within the same cell. Given the extremely low copy number of mRNA in the periphery of axons and dendrites and their small diameter, current spatial transcriptomics technologies such as Nanostring, lack both resolution and sensitivity for systematic spatial characterisation of transcriptomes in intact nervous systems. Moreover, single cell transcriptomics approaches lose the peripheral compartments of cells, so are not applicable to systematically address peripheral localisation in neurons. Single molecule FISH methods such as merFISH can overcome the issues of resolution and sensitivity, but are not compatible with systematic spatial protein analysis.

Here, we have overcome these technical limitations by developing a widely applicable workflow for comparing the level of discordance between hundreds of mRNAs and their corresponding proteins at high resolution across complex tissues in 3D. Our approach depends on the use of a fluorescent protein to tag many individual endogenous genes, and systematic visualisation of mRNA using smFISH to detect greater than 50% of all individual tagged molecules of mRNA in every cell at high resolution. mRNA detection is coupled with co-visualisation of protein at high sensitivity and resolution in the same specimens. We have prototyped our workflow for 200 genes in the intact nervous system and neuromuscular junction of third instar *Drosophila* larva using a collection of YFP fusions (Lowe et al. 2014). Our unexpected results led us to a wholesale revision of the global view of post-transcriptional regulation, mRNA localisation and delivery of proteins to the periphery of the nervous system. Post-transcriptional regulation is very common across all of the nervous system, acting hand in hand with transcriptional regulation to create a complex tapestry of protein distribution in time and space. We present our data as a resource that is easily browsable in the context of a rich landscape of genomics, functional and bioinformatics data, using Zegami (Taylor and Noble 2014), a web-browser based software for interactive data visualisation and exploration.

## Results

### Systematic analysis of the level and distribution of the mRNAs of 200 genes by smFISH against YFP fusions and their corresponding fluorescent protein visualisation across the nervous system

To ask how gene expression is controlled in specific cell types and subcellular compartments in the nervous system, we developed an imaging pipeline to simultaneously quantify transcription, mRNA and protein levels throughout whole tissues for hundreds of different genes (Figure 1A). Our scalable approach takes advantage of *Drosophila* gene trap collections that have a fluorescent protein reporter inserted into introns of individual genes, flanked with splice donor and acceptor sites. Using a common single molecule fluorescence *in situ* hybridization (smFISH) probe against the mRNA sequence encoding yellow fluorescent protein (YFP), we detected reporter mRNAs along with an encoded reporter YFP protein. The smFISH probe also acts as a transcription reporter by detecting primary transcripts at the endogenous gene locus in nuclei. Imaging the smFISH probe and fluorescent protein tag in whole tissues with confocal microscopy allowed us to systematically map the spatial distribution of gene expression in many different regions and cells of the nervous system at high sensitivity and resolution.

**Figure 1.**
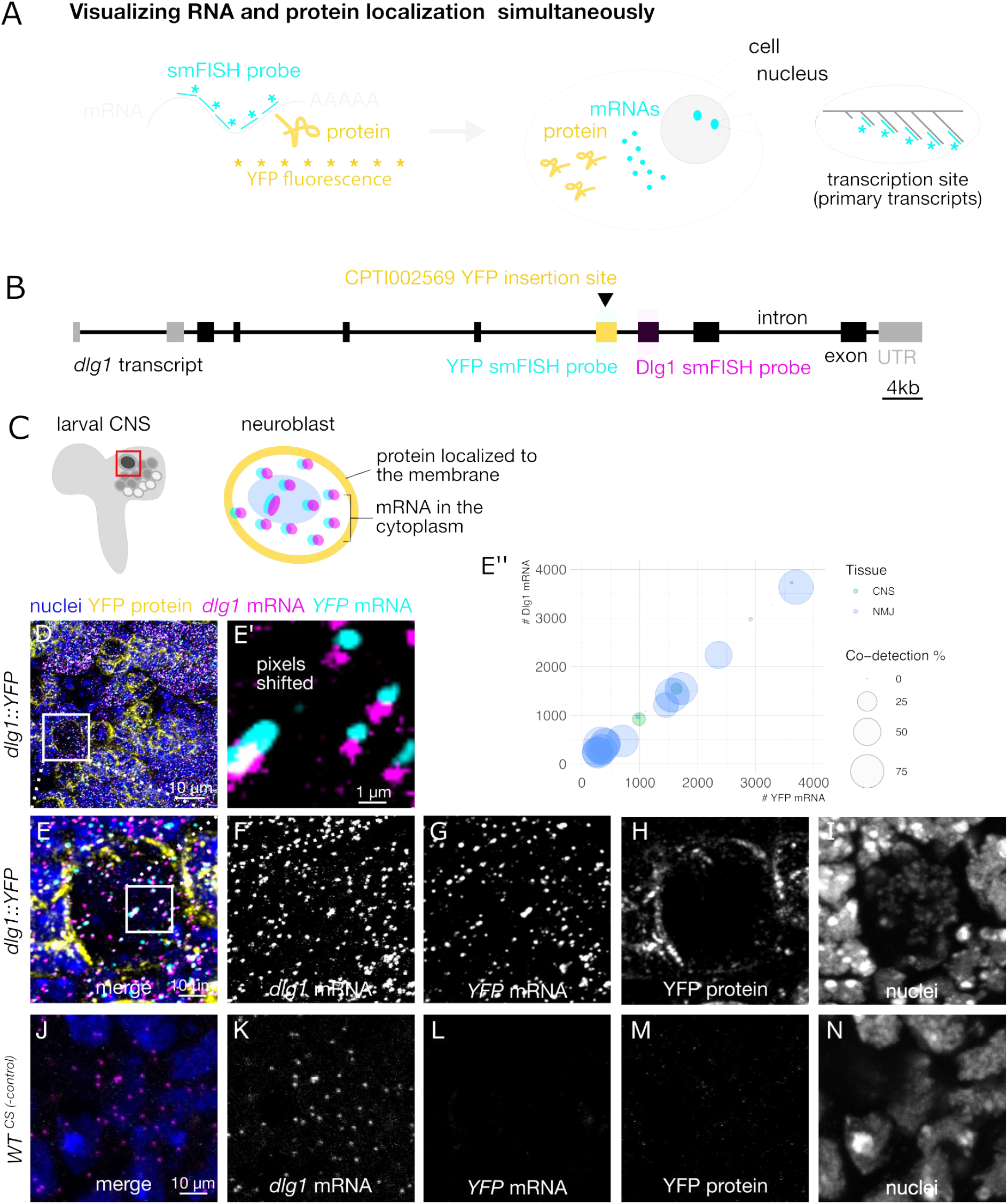
Spatial detection of localised mRNA and protein expression across multiple tissues and hundreds of different genes in *Drosophila* larvae. (A) General strategy for simultaneously visualising RNA and protein in fluorescent protein trap lines. RNA is detected using smFISH probes targeting the genetically encoded yellow fluorescent protein (YFP) sequence that is present within the mRNA expressed from the trapped gene. Large transcription foci are seen at the gene locus where there are multiple primary transcripts. Protein is detected by fluorescence of YFP in the protein. This approach can be used to detect any trapped gene in any tissue. Here we have focused on the nervous system. (B) Genetic architecture of CPTI002569, a YFP gene trap line that was used for one set of control experiments. This line has a YFP reporter inserted into an intron that is contained in the *dlg1* transcript. smFISH probes were designed to target the YFP sequence or a common *dlg1* exon. The probes were labelled with spectrally separated dyes to perform a co-detection experiment. (C) Schematic showing a region of the *Drosophila* larval central nervous system that was imaged to visualise *dlg1* expression in neuroblast lineages. The inset schematic shows the endogenous expression pattern of *dlg1* mRNA and protein in a single neuroblast. (D) Overview maximum intensity projection confocal image from a *dlg1::YFP* line showing expression of YFP protein (yellow), YFP smFISH probe (cyan) and *dlg1* smFISH probe (magenta). Signal from both smFISH probes was observed as individual diffraction-limited punctae throughout the larval central brain. (E) Positive control experiment-high magnification image of the inset in D showing individual transcripts and protein expression within a single cell. E’-inset, YFP signal was intentionally shifted by 3 pixels to visualise transcripts that were detected by smFISH probes against two different sequences within the *dlg::YFP* transcript. (E’’) Percentage of *yfp* and *dlg1* co-detection in both CNS and NMJ tissues. (F-I) Grayscale images of the individual channels shown in E. (J) Negative control experiment-maximum intensity projection image of the same region in a wild type line that does express YFP. Note that *dlg1* transcripts are detected by the *dlg1* smFISH probes, but there is no signal in the YFP smFISH channel. (K-N) Grayscale images of the individual channels shown in E.

As proof-of-principle, we performed smFISH experiments on a Discs large 1 protein trap line (Dlg1::YFP) in the larval central brain (Figure 1B-C). *Dlg1* (PSD95 in mammals) is a tumour suppressor gene encoding a protein that localises to intercellular junctions (Peng et al. 2000; Albertson and Doe 2003). We found that our YFP smFISH probe is highly sensitive and specific for *dlg1::YFP* mRNA, and that the reporter insertion does not affect the localisation of the *dlg1* mRNA or protein. To determine the specificity of the YFP smFISH probe we tested whether the probe detects any transcripts in a wild type line that lacks YFP (Figure 1). While in the *dlg1::YFP* gene trap line, the YFP smFISH probe labels hundreds of diffraction-limited punctae throughout the central brain (Figure 1D-I), no equivalent signal could be detected in wild type samples (Figure 1J-N). The majority of individual puncta appearing in the *dlg1::YFP* line (51% in the brain, 64% in larval muscles (Figure 1E’’)) were also detected by a spectrally separated second oligonucleotide probe set targeting the endogenous *dlg1* transcript, indicating that the probe is highly sensitive (Figure 1D). Importantly, the Dlg1::YFP protein showed its characteristic enrichment at the cell surface, which means that the reporter protein does not disrupt localisation or expression level of the endogenous protein. We conclude that this is an effective approach to screen for gene expression patterns and proceeded to apply the method to 200 gene insertions randomly selected from the Cambridge Protein Trap Insertion (CPTI) collection (Lowe et al. 2014). We not only imaged the central brain and neuroblasts (neural stem cells), but also the mushroom body (equivalent to mammalian hippocampus), optic lobe, ventral nerve cord, segmental nerves, and the larval neuromuscular junction neurons, muscles and associated glia.

To determine how well the selected set of genes captures the diversity of gene expression patterns in the whole transcriptome, we analysed published data on the gene expression levels, gene structure, and gene functions. We found that this set of 200 genes provides a fairly representative sample of transcript heterogeneity. We first analysed publicly available bulk RNAseq data from specific tissues and developmental stages. Given that CPTI lines were selected for the presence of YFP reporter expression in embryos, we compared the overall distribution of gene expression levels in embryos to third instar larval brains. Violin plots show that the distributions of gene expression levels are similar in the two tissues, and by overlaying the individual genes that were screened in the current study (Figure S1A), it is clear that the genes we analysed are relatively abundant and span the entire range of gene expression levels, making the collection a useful proxy for the whole transcriptome.

Next, we characterised the physical structure of the 200 screened genes. The CPTI collection was created by a hybrid *piggyBac* vector insertions which favours longer genes and longer introns, since the gene traps are formed by random insertions into introns (Lowe et al. 2014). We found that the 200 CPTI genes we analysed are indeed as expected on average slightly larger, and contain longer introns than the average protein coding gene (Figure S1B). Since it is thought that genes that are highly expressed in the nervous system tend to be longer and contain longer introns than average (McCoy and Fire 2020), we conclude that our 200 genes are likely to be enriched in genes that are highly expressed in the nervous system.

In contrast, we found that the 3’UTR extension lengths were similar in the 200 CPTI lines compared with the average protein coding gene (Figure S1B). Given that the majority of known localisation signals reside in 3’UTRs (Tushev et al. 2018), we interpret our mRNA localisation results as being representative of the whole genome. Similarly, 3’UTRs extensions often contain sequences that regulate mRNA stability, suggesting that the 200 CPTI genes are likely to be similar to the rest of the genome, at least in the characteristics of their 3’UTR extensions.

To assess how representative the 200 genes in our screen are for gene function in the genome, we compared the total number of unique parent GO terms (GOSlim terms) associated with the genes in our dataset to the number of unique GO terms found in all protein coding genes. The genes in our dataset map to 89.9% of the terms across all three GO categories (The GO categories are available in Supplementary Table S1), which makes the collection highly representative of the functional diversity of protein coding genes. Together, these results indicate that we are not significantly undersampling the complexity of gene expression patterns, and that the percentage of genes with a given expression pattern in our sample could be extrapolated to provide an estimate of the total number of transcripts with that expression pattern across the whole transcriptome.

### A generalisable workflow for assembling and browsing integrated microscopy and bioinformatics databases

Extracting biological insight from large microscopy datasets is a notoriously challenging and laborious process. To facilitate analysis and browsing of our dataset we established a generalisable workflow (Figure 2) to display the images, annotate and score gene expression patterns consistently across many cell types, and systematically interrogate the microscopy data together with genomics data and other large scale microscopy studies. This approach makes the data easier to interpret, facilitates novel insight and hypothesis generation, and extends the functionality and utility of published resources.

**Figure 2.**
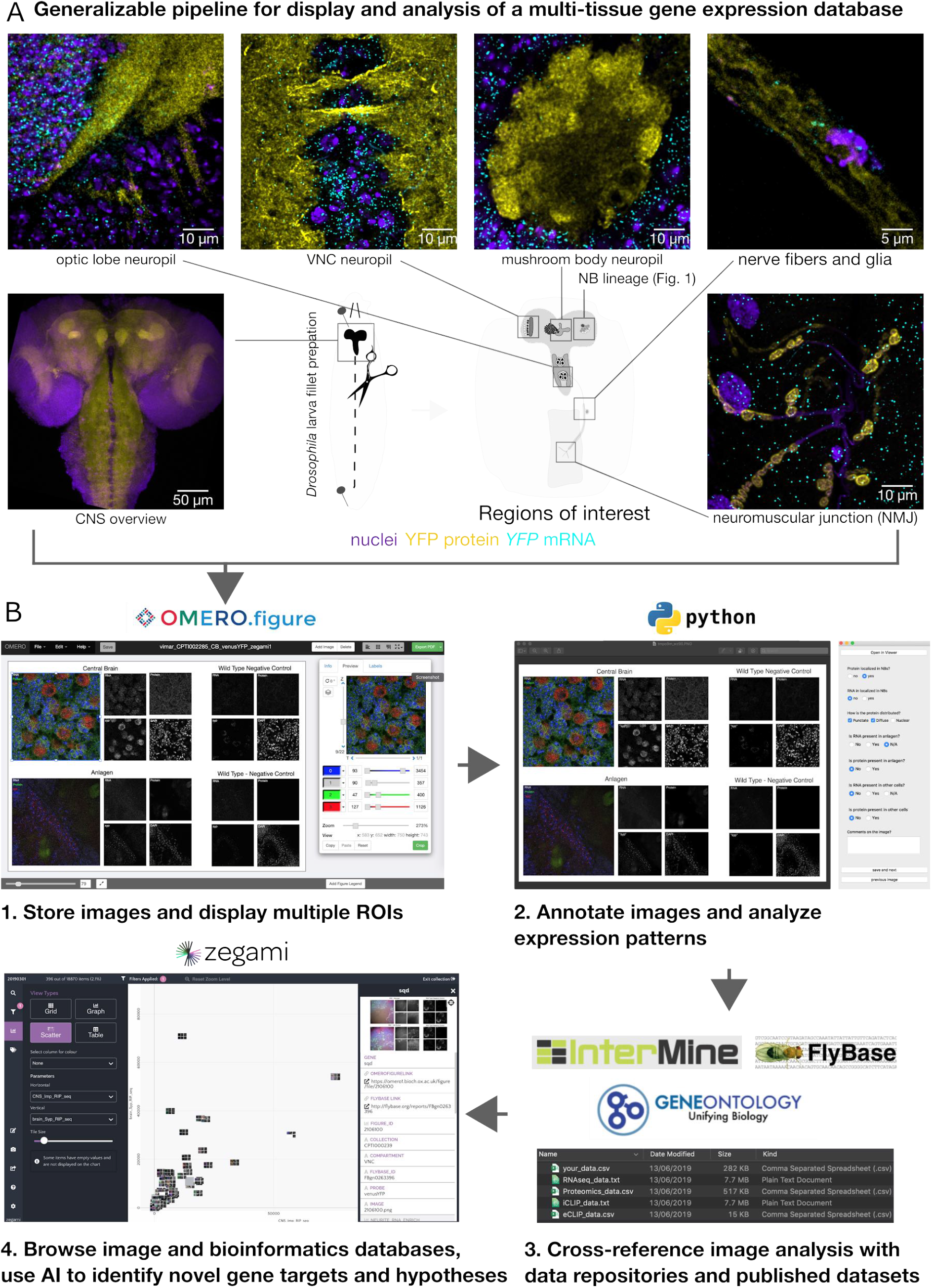
A custom image annotation application and generalizable workflow to assemble and browse integrated imaging and bioinformatics databases. (A) Images are obtained from multiple nervous system compartments. 1) Microscopy data is stored on an OMERO server and adjusted for multidimensional display using the OMERO.Figure web application. 2) A customizable user interface was developed in Python to display and annotate the OMERO.figure images. 3) A Python application with graphical user interface is used to write annotations to a database along with queries from publicly available bioinformatics datasets. 4) The database can then be imported into the Zegami web application to intuitively explore the data and discover hidden functional associations with machine learning algorithms.

The images in our dataset, like most light and electron microscopy images, contain rich and diverse 3D information that is difficult to convey in a single snapshot, or a single figure in a manuscript. Each image consists of a large 3D volume in which there are multiple cell layers and multiple labels that can be used to address different biological questions throughout the volume. Moreover, each gene was characterised in multiple tissues of the nervous system, so the combined dataset represents more than 1000 individual figures, a data volume that cannot be published as conventional figures in a manuscript. Therefore, we developed an approach that displays selected views of the 3D image stacks simultaneously, while also providing access to the raw intensity data. Using the open source OMERO.Figure web application (Allan et al. 2012) with its links to the original data stored in OMERO (Goldberg et al. 2005). Multiple regions of interest (ROIs) from specific compartments were selected and contrasted to display specific cellular compartments from each image in an easily browsable and consistent ‘Figure’ format, at scale.

To also analyse figures quantitatively at scale, we developed a Python application to systematically annotate OMERO.Figure images, which we named Annotate.OMERO.Fig (see Materials and Methods). The scoring application takes a customisable set of questions, which are presented to scorers via a graphical user interface as it cycles through an image dataset. Then, the user-scored answers are collated and exported in a spreadsheet format for downstream analysis. Three experts annotated each tissue independently by answering the same standardised questions, such as “is RNA present in axon terminal?”. Where the expert annotators disagreed, we used a majority vote approach to select the correct answer (Figure 3). Since it was important to also view the results in the context of what is already known about a gene, we added a script to extract data from specified databases, either from online repositories or directly from local files, and merge the data into a single file. This approach allowed us to browse images associated with a specific gene, while simultaneously viewing its gene ontology, relative expression levels in published transcriptomic and proteomic datasets, and genetic screens. Moreover, assembling the imaging data in its rich bioinformatic meta-data context made it possible and convenient to deploy machine learning algorithms for hypothesis generation and gene candidate selection to guide future experiments.

**Figure 3:**
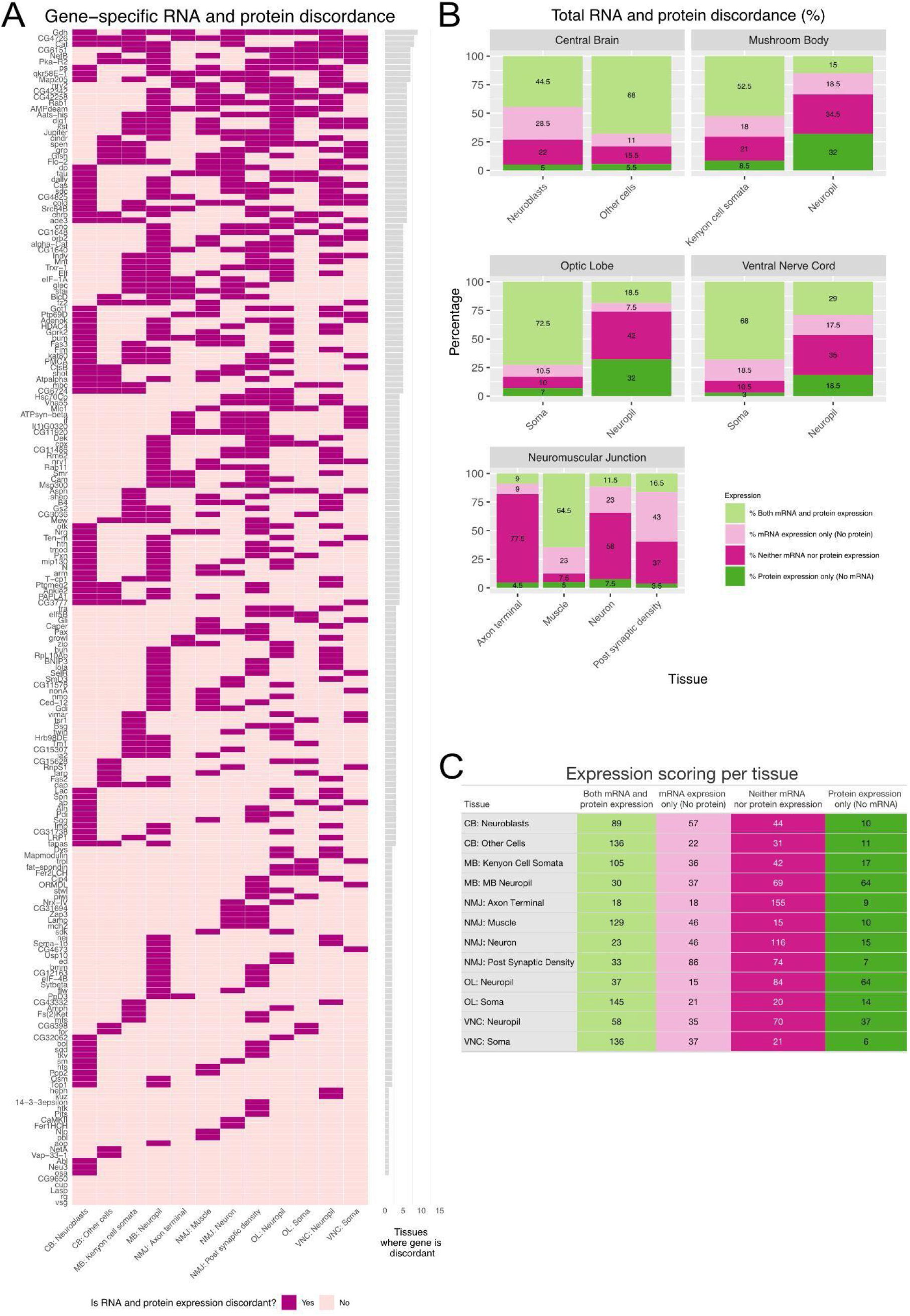
Summary of annotations for mRNA and protein expression for each tissue and discordance. (A) Discordance per gene per tissue shown in dark purple and concordance in light pink. Each row corresponds to a gene and the reader can see how many tissues show discordance per gene. (B) Percentage of genes scored and (C) numbers of genes scored that show expression of mRNA and protein or the absence of expression of mRNA and protein per tissue and compartment. The graph also shows percentages of mRNA expressed and protein not expressed and vice versa per tissue and compartment.

To facilitate manual curation and data browsing, we uploaded the annotation file and associated images into a web application called Zegami, an AI-enabled image analysis platform used to gain high level insight from large datasets. We designed and built a pipeline that is easily generalisable to other model organisms and data repositories. Our image dataset includes 1,361 Figures from 200 genes in the CPTI gene trap collection, with downstream analysis of the whole genome, allowing for extrapolation of the findings to predict additional genes with similar expression patterns or phenotypes. Moreover, the dataset lists other genes with known protein trap insertions and links directly to Intermine (Smith et al. 2012) and FlyBase (Larkin et al. 2021), which extends the utility of those resources.

### Overview of the screen results

A fundamental question we addressed with this dataset is where proteins and mRNAs are expressed relative to each other throughout different cell types of the nervous system for each of the 200 genes (see tables in Figure 3 for a visual summary of scoring results for each gene in each compartment). Below, we describe our findings of the quantitation of mRNA and protein correlation in each compartment. Compiling all the information together shows that there are that 196/200 or 98% of the genes show discordance between RNA and protein expression in at least one cellular compartment of the nervous system, and 137/200 or 68.5% of genes show RNA localisation in at least one of the synaptic regions we examined (Supplementary Table S2). Our dataset represents a unique and rich resource describing the discordance between mRNA and protein for each gene in each compartment at high 3D spatial resolution and very high sensitivity of single molecule mRNA detection (Figure 3A). The data set can also be interrogated in a complementary way to discover the percentage (Figure 3B) and total number of genes (Figure 3C) that express RNA and/or protein in each compartment in the larval nervous system.

### Post-transcriptional control of neuroblast differentiation

Although transcription factors have been thought to be the primary regulators of neuronal differentiation, post-transcriptional regulation also plays a major role in nervous system biology (Cajigas et al. 2012). To assess the prevalence of post-transcriptional regulation in neuronal differentiation, we applied our mRNA and protein reporter approach to visualise gene expression in populations of asymmetrically dividing neuroblasts in the larval central brain (Figure 4), a powerful and well used model for understanding neural differentiation (Homem and Knoblich 2012). We discovered that post-transcriptional regulation is unexpectedly widespread among genes that are selectively expressed in neuroblast lineages (see Supplementary Table S3 for detailed analysis associated with Figure 4). Approximately one third of the genes with cell-specific expression in our dataset show discordance between protein and mRNA expression in neuroblast lineages (Figure 4A), a hallmark of post-transcriptional regulation. Cell-specific expression patterns were observed for 21.5% of the genes (43 genes), whereas 57% of the genes (114 genes) were expressed homogeneously throughout the neuroblast lineage (Figure 4B), and 21.5% of genes (43 genes) were not detected above background at either the mRNA or protein level (Figure 4C). We found that every gene that expresses mRNA or protein in neuroblasts is also expressed in their immediate progeny, indicating that none of the 200 genes in this dataset are strictly neuroblast-specific in their expression. In fact, most of these genes are expressed broadly throughout the central brain. However, one gene, *indy*, is highly transcribed in neuroblasts and a single ganglion mother cell before it is rapidly shut off (Figure S1A). We conclude that post-transcriptional regulation is likely to play a wide-spread role in neuroblast biology and differentiation of its progeny.

**Figure 4.**
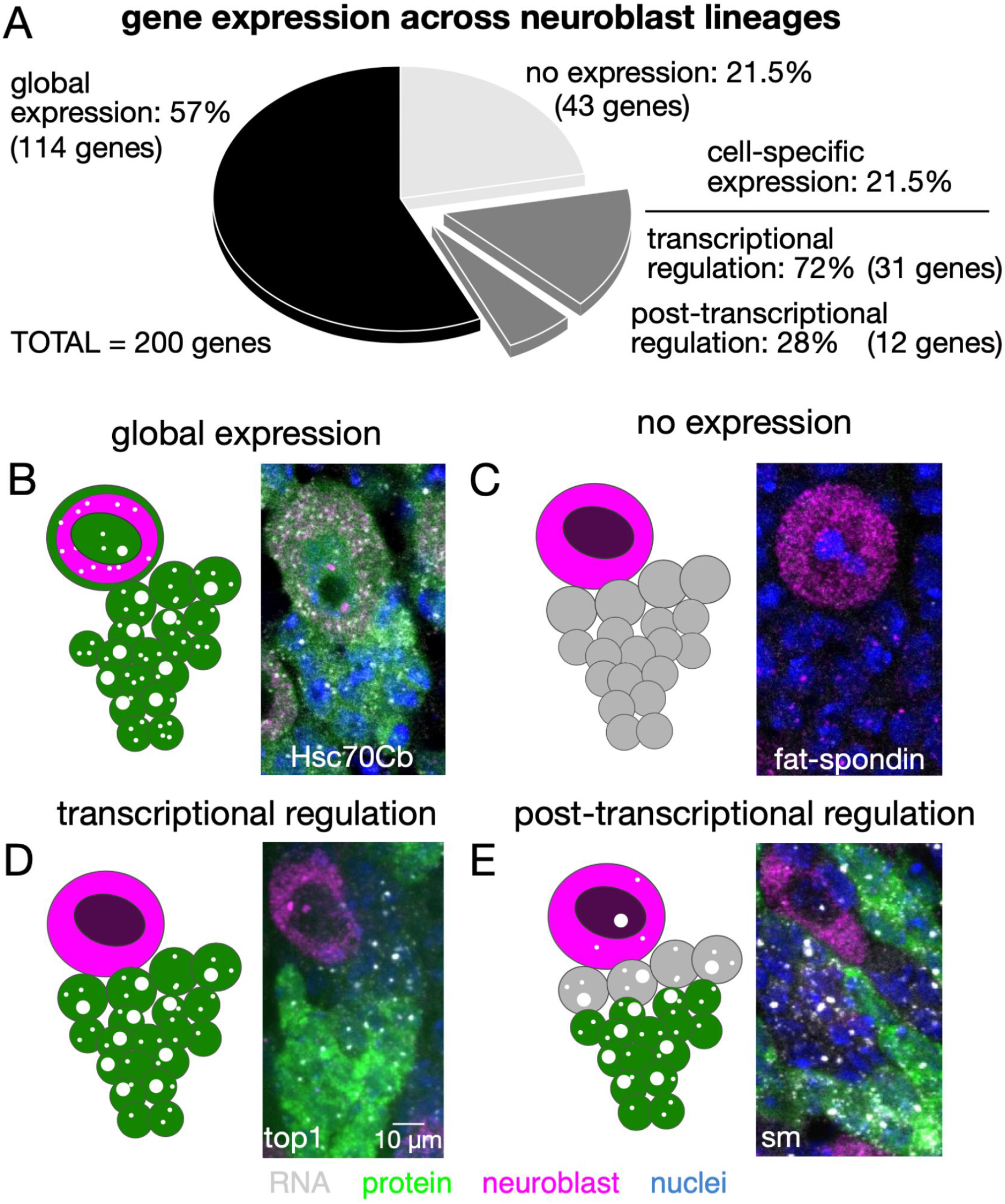
Discordance between protein and mRNA expression patterns reveals the extent of post-transcriptional regulation in neuroblast differentiation. Two hundred YFP-reporter lines were imaged across multiple type-I neuroblast lineages in the larval central brain. (A) Pie chart showing the relative distribution of different gene expression patterns. (B) The majority (57%) of genes were expressed homogeneously throughout the neuroblast lineage, while 22% of genes were not detected in the central brain region (C). The remaining 21% of genes exhibited some degree of cell-specific expression throughout the neuroblast lineage. In some of the 21%, where protein levels were either correlated with mRNA and transcription levels indicating transcriptional regulation (D), or not correlated with mRNA levels indicating post-transcriptional regulation (E). Over one quarter of all genes with cell-specific expression patterns exhibited this hallmark of post-transcriptional regulation.

Of the 43 genes with cell-specific expression patterns, 31 (72%) genes exhibit highly correlated protein and mRNA expression between different cell types. Of those 31 genes, only a subset of cells actively transcribes the gene, and each cell that produces mRNA also produces protein. Highly correlated protein and mRNA expression is a strong indication that these genes are transcriptionally regulated. A representative gene with this expression phenotype is *top1*, a topoisomerase that has essential functions in cell proliferation (Figure 4D). Of the 43 genes with cell-specific expression, 12 (28%) exhibit obvious discordance between mRNA and protein levels throughout the neuroblast lineage (Figure 4A). The transcription rate of these genes, as indicated by the relative intensity of smFISH nuclear transcription foci, is similar across the neuroblast lineage, however protein signal is only detectable in a minority of the progeny cells (Figure 4E). Discordance between protein and mRNA content is a strong indication of post-transcriptional regulation, and also suggests an important cell-specific function. Consistent with this idea, four of the twelve genes with discordant expression patterns, *pbl, Rm62, qkr58E-1* and *cno*, were previously identified in a genome-wide screen surveying neuroblast division phenotypes (Neumüller et al. 2011).

### Synaptic mRNA localisation across different CNS neuropils

mRNA localisation provides an additional layer of post-transcriptional regulation to target specific proteins to neural synapses. To determine the contribution of mRNA localisation to synaptic proteomes we visualised mRNA and protein content across multiple neuropil regions of the intact larval brain. We found that nearly half of the genes in our dataset express proteins that are relatively abundant at mushroom body synaptic regions, at the periphery of cells. Nearly one third of the genes also express mRNA that is present at the synapse. However, the sets of synaptic mRNAs and synaptic proteins do not overlap entirely, providing insight into the specific mechanisms of localised post-transcriptional regulation (see Supplementary Table S2 for detailed analysis).

To analyse these localisation patterns further, we acquired stacks of confocal images from three synaptic regions in the larval nervous system, the mushroom body, the optic lobe neuropil, and the sensorimotor neuropil of the ventral nerve cord. For each of the 200 randomly selected YFP protein trap lines, we assessed whether the protein and/or mRNA was expressed in soma or in synaptic neuropil for each region of interest. In the mushroom body, 94 out of the 200 genes in our dataset (47%) encode proteins that are detectable either in the mushroom body calyx or one of the lobes. Of those 94 genes, 30 (32%) are encoded by mRNAs that are also detected in either region of the mushroom body (figure 5A-D). These observations suggest that localised mRNA translation contributes about one third of the synaptic proteome, slightly less than what has been previously reported (Zappulo et al. 2017). Surprisingly, another 59 transcripts are present at synapses without detectable levels of protein (Figure 5E-H), and unexpectedly, many of those genes encode proteins with predominantly nuclear localisation and function (Figure 5I-L, Q). We hypothesise that these mRNAs are translationally silenced and primed to produce proteins that will transduce synaptic input to the nucleus. However, we cannot exclude the possibility that these mRNAs encode proteins that are present at levels below our detection limit, or that these transcripts are present in neuropil, without performing a function.

**Figure 5.**
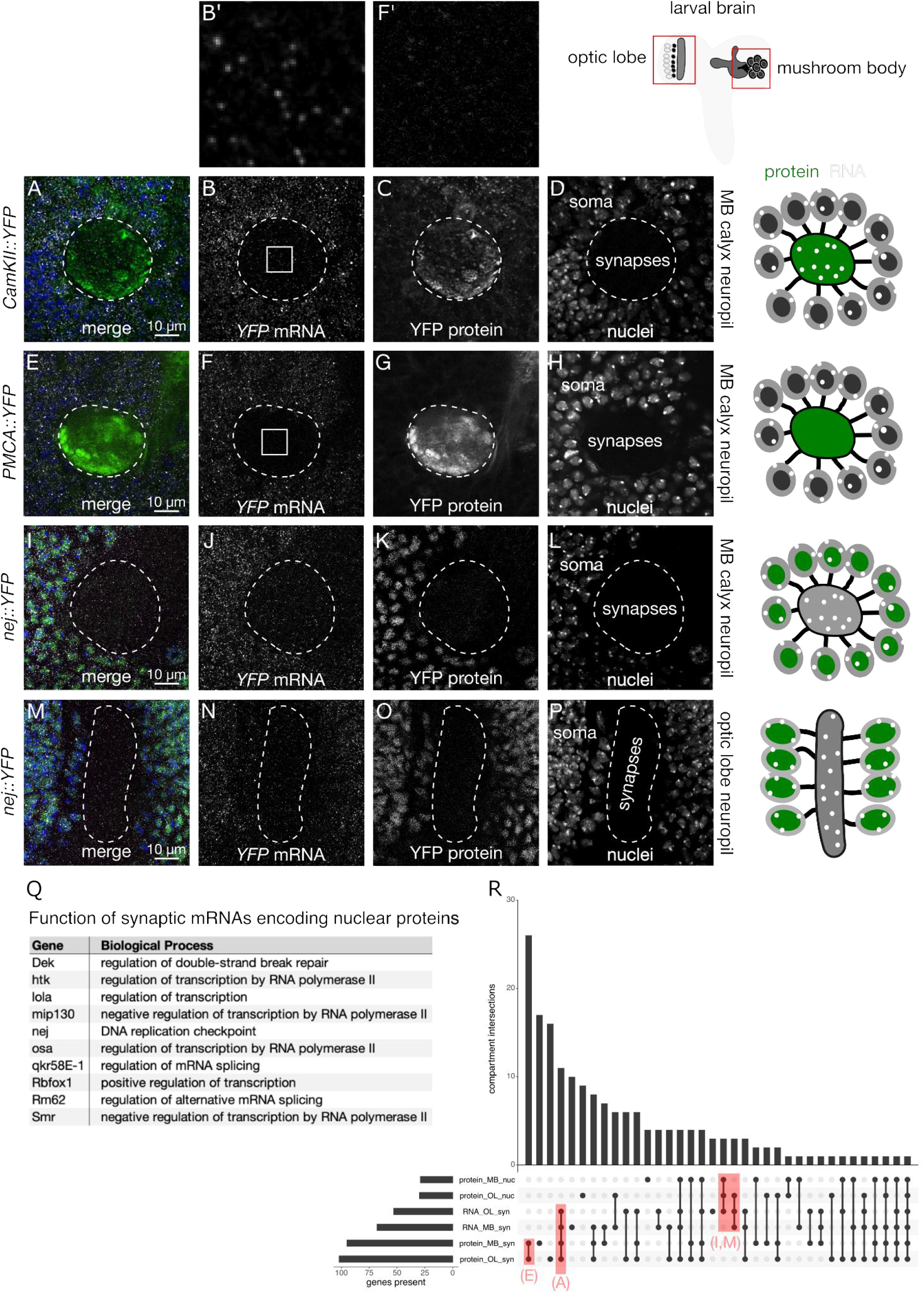
Discordance between synaptic localisation of mRNA, protein, and synaptic function is surprisingly prevalent. (A-H) Optical sections of the larval brain showing mRNA and protein distribution patterns in the mushroom body (MB) and optic lobe (OL) neuropils. The majority of synaptic proteins have mRNAs at the synapse (A-D), but 47% of synaptic proteins are expressed without localised mRNAs (E-H). Nearly half of the synaptic mRNAs fail to generate detectable levels of synaptic protein, instead, these mRNAs tend to encode nuclear proteins in both the MB (I-L) and OL (M-P) neuropils. Surprisingly, many of the proteins encoded by synaptic mRNAs are transcription factors (Q). (R) UpSet plot showing the number of genes expressed in each compartment, at the mRNA and protein level, for the entire dataset.

We reasoned that it would be unlikely for synaptic localisation of mRNA that lacks any function to occur consistently across different cell types. Therefore, we repeated the localisation analyses in different types of neurons, including the optic lobe and VNC neuropils. We found that 28 of the 67 (42%) mRNAs present at the mushroom body synaptic neuropil are also present at the optic lobe neuropil. Moreover, many of the synaptic mRNAs that encode nuclear proteins were also present in the optic lobe neuropil (Figure 5M-P).

### mRNA and protein localisation in glia

Like neurons, many glial cell types have long and elaborate filamentous processes that are likely to require localised gene expression control through mRNA localisation and local translation. Though localisation of mRNAs encoding glial fibrillary acidic protein and myelin basic protein have been extensively characterised (Medrano and Steward 2001; Müller et al. 2013), relatively little is known about the hundreds of other localised mRNA species that have recently been identified in mammalian glial processes, and almost nothing is known about mRNA localisation in *Drosophila* glia.

To identify new genes with mRNA localised to glial processes in the larval central nervous system we manually searched the image dataset for protein and mRNA expression patterns that show enrichment in glial processes. There are six types of glia that have invariant positions and characteristic morphologies within the nervous system (Schmidt et al. 2012). We focused on cortical glia in the central brain region, and ensheathing glia that envelop the mushroom body neuropil, as the processes in these cells are easily identifiable. For reference, we labelled those regions by expressing membrane-bound RFP specifically in glial cells using the repo-Gal4 driver (Figure 6A-B,E).

**Figure 6.**
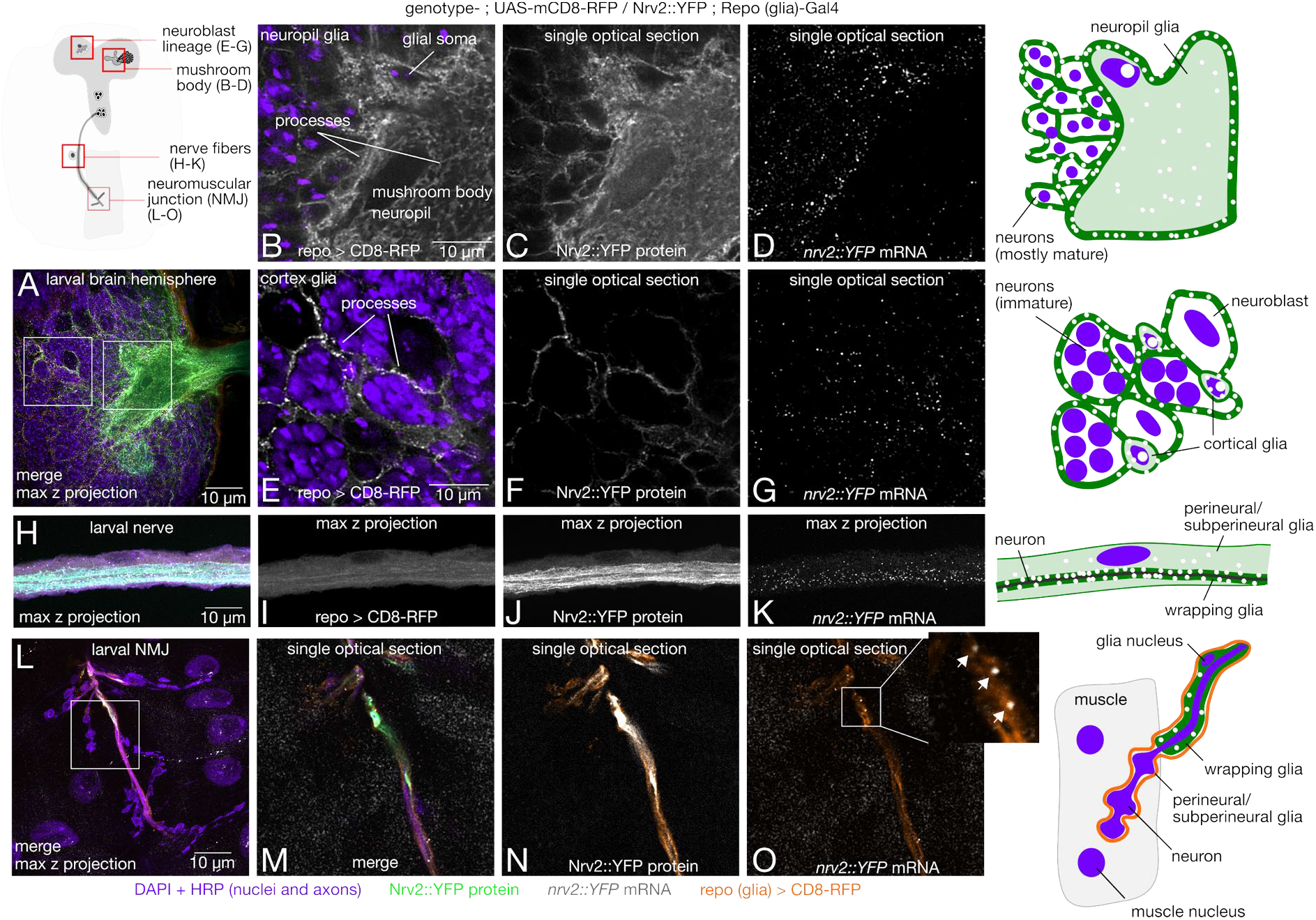
mRNA is localised in glial processes throughout the larval nervous system. Repo (glia)-Gal4 and UAS-mCD8-RFP (membrane marker) were crossed into the Nrv2::YFP background to demonstrate *nrv2* mRNA (grey) and protein (green) localisation in glial processes (orange) throughout the larval nervous system. (A) Overview confocal maximum intensity projection image of a larval brain hemisphere showing the relative locations of glial cells. (B-G) Single optical sections showing spatial overlap of *nrv2::YFP* RNA and protein channels with the repo > RFP marker in neuropil (B-D) and cortical (E-G) glia. (H-K) Representative image of *nrv2::YFP* mRNA and protein expression in segmental nerves innervating the larval body wall musculature. (L) Overview of glia anatomy and *nrv2::YFP* expression at the larval neuromuscular junction (NMJ). (M-O) Single optical sections showing spatial overlap between glial processes, Nrv2::YFP protein and single mRNA molecules (inset, arrows).

To assess glial localisation for the 200 genes of interest, we used a pan-glial gal4 driving a membrane mCherry marker (*repo-GAL4>UAS-mcd8-mCherry*) to learn the expression pattern of all glial cells, and then classified the pattern in the YFP lines (without the marker) based on knowledge of that expression pattern. We validated this approach by combining the RFP marker with the *Nrv2::YFP* insertion, one of the 200 lines which was already known to be a wrapping glial marker (Yadav et al. 2019). We performed smFISH using probes against the YFP sequence and found that *Nrv2* protein and mRNA are highly expressed in glial processes of both cell types (Figure 6C-D, F-G) demonstrating that we can accurately classify glial mRNA localization based on the stereotypical glial morphology observed in the protein expression pattern.

Focusing on cortical and ensheathing glia in the central nervous system, we found that 19.5% of the proteins in our dataset are expressed in cortex glial processes in the central brain, and only 11.5% of proteins are expressed in mushroom body neuropil glial processes. A very high percentage of mRNAs that encode those proteins are also localised to glial processes, 92% and 65% for the cortex and neuropil glia respectively (Supplementary Table S3).

We extended our analysis of glial mRNA localisation to glia in peripheral nerve fibres and at the neuromuscular synapse. Perineurial and subperineurial glia wrap the outer layers of the nerve bundle, and wrapping glia envelope single axon fibres. The glia also form the blood nerve barrier between the axon and extracellular fluid. Perineurial and subperineurial processes extend into the neuromuscular synapse. Each of these cell types is marked by the repo > mCherry reporter along the fibre (Figure 6H-I) and at the neuromuscular synapse (Figure 6L-O). *Nrv2* mRNA and protein are distributed throughout each layer of glia in both the nerve fibre (Figure 6J-K) and axon terminals (Figure 6N-O) with highest expression in the wrapping glia, which form the blood nerve barrier between the axon and extracellular fluid. We also show that mRNAs, e.g., *gli*, are localised in the subperineurial and perineurial glia that are associated with more distal boutons (Figure 6S). Focusing on axon terminals at the larval NMJ, where glial protein expression patterns are easily identifiable, we found 19 genes (9.5%) with protein or mRNA expression in either wrapping glia at axon terminal, or perineurial and subperineurial glia of the NMJ, and 95% of those genes also showed mRNA localisation in the glial processes. Together, analysis of mRNA and protein expression in glial processes of the CNS and PNS shows that mRNA localisation makes a major contribution to the proteome in that compartment (Supplementary Table S3). Our results present the first examples of mRNA localisation to glial processes in the larval central nervous system, segmental nerve, and neuromuscular junction, highlighting a hitherto unrecognised important general function for mRNA localisation and most likely, localised translation.

### mRNA and protein localisation at neuromuscular synapses

mRNA localisation to motor axon terminals is one of the most extreme examples of polarised gene expression in metazoans, with mRNAs being transported, within some neurons, a distance nearly equivalent to the entire body length of the animal. Neuromuscular synapses on the larval body wall muscles provide an excellent system to investigate such long distance axonal mRNA transport, to determine whether the mRNAs are pre or postsynaptic, a question that is much harder to address in the dense synaptic neuropils of the central brain. To investigate how frequently different mRNAs are localised, we applied our mRNA and protein trap microscopy screening approach to the larval neuromuscular junction. We found that the presence of mRNA in these motor axon terminals is relatively rare, as is strong enrichment of mRNA in the postsynaptic density (PSD) in the muscle cells.

We combined our smFISH and protein trap approach with subcellular markers to distinguish individual axon terminals, the postsynaptic density and muscle nuclei. We found that 13.5% of the genes in our dataset encode proteins that are detectable in the motor axon terminal (Supplementary Table S2). Around two thirds of those proteins are accompanied in the axon terminal by the mRNAs that encode them (Supplementary Table S2), which is consistent with the percentage of transcripts that are typically detected in transcriptome-wide studies (von Kügelgen and Chekulaeva 2020). An example of a gene with this expression pattern is *sgg* (Figure 7A-D), which we chose to further characterise because of its known role in NMJ development (Figure 8). We only observe a minority of localised axonal mRNAs that lack the protein they encode at the axon extremities, in contrast to our findings in the mushroom body, optic lobe, and ventral nerve cord neuropils. These results suggest that motor axons are more selective than the other neuronal extensions in the mRNAs that are transported over their very long distances from the soma to the neuromuscular synapse.

**Figure 7.**
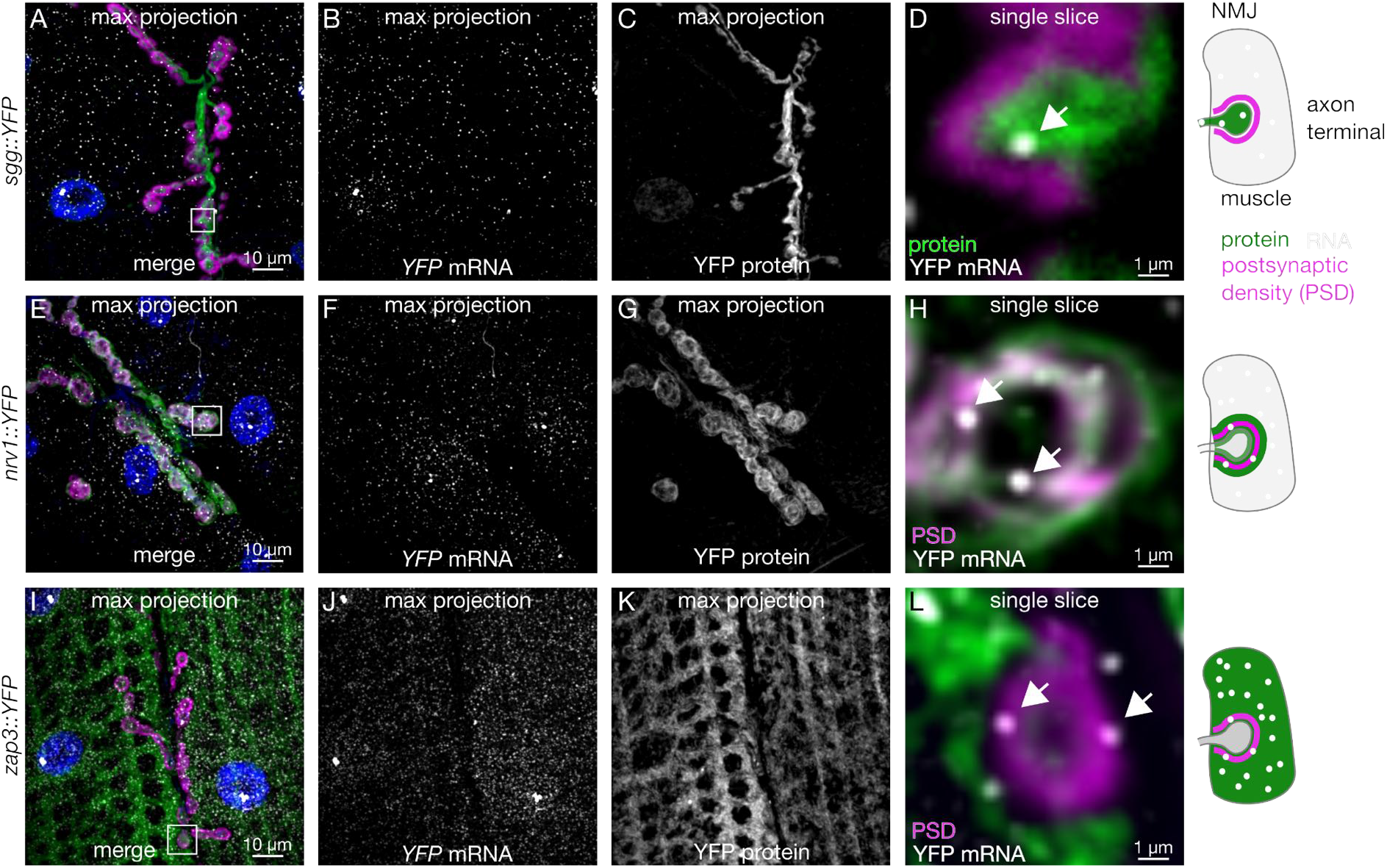
mRNA is present on both sides of the larval neuromuscular synapse. (A-C) Maximum intensity projection of confocal images showing *sgg::YFP* mRNA and protein localisation at the larval NMJ. (D) Single optical section of the region in A (white box, 10x magnification) showing protein and individual mRNA molecules (arrow heads) located in the axon terminal. (E-G and I-K) Max projections of *nrv1::YFP* and *zap3::YFP* expression, which have very distinct protein expression patterns despite nearly indistinguishable mRNA patterns. (H and L) High magnification single optical sections show that mRNA from both genes are present within the postsynaptic density.

**Figure 8.**
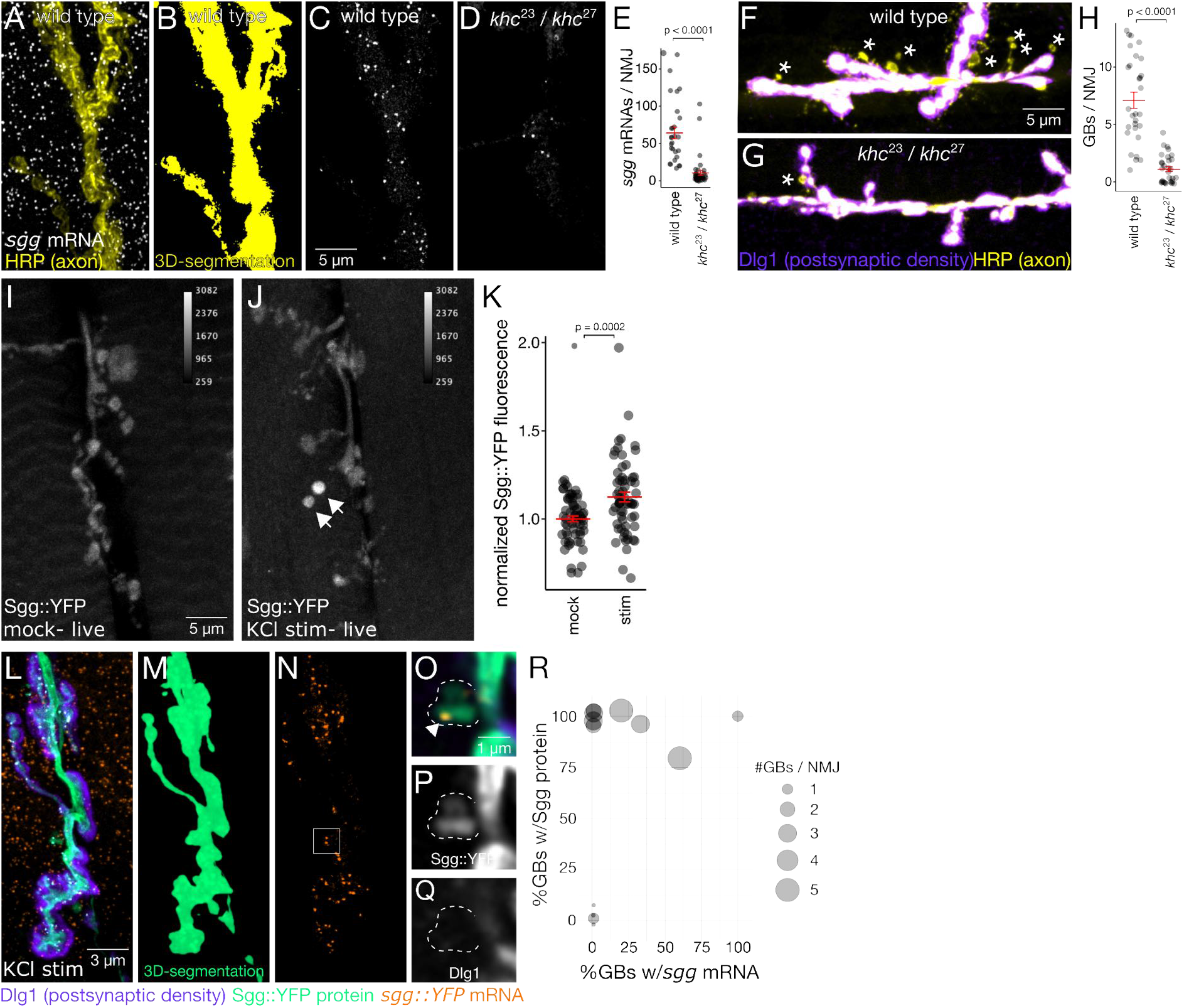
*sgg* mRNA is actively regulated at the larval neuromuscular junction. (A-E) *sgg* mRNA localisation at axon terminals requires kinesin. (A) Maximum z-projection image showing *sgg* smFISH signal at the axon terminal (yellow). Images were segmented in 3D with the axon marker channel (B), revealing a significant decrease in the number of axonal *sgg* mRNAs in *khc*^*23*^*/khc*^*27*^ transheterozygous mutants (C-D). (E) Quantification of *sgg* transcript levels in axon terminals (mean ± SEM; Student’s t-test). (F-H) Kinesin is required for ghost bouton formation. Max z-projection images show GBs (asterices) in KCl stimulated NMJs from kinesin mutants and wild type controls. (H) Quantification shows significantly fewer GBs in kinesin mutants (mean ± SEM; Student’s t-test). (I-K) Live imaging of axon terminals shows a significant increase (mean ± SEM; Student’s t-test) in Sgg protein levels in samples stimulated with five pulses of KCl. (L-R) In fixed samples, Sgg protein and mRNA are present in a large percentage of activity-induced synaptic boutons (ghost boutons-GBs). Axon terminals were segmented in 3D to isolate signal in the axon from signal in the muscle (M). (O-Q) Single optical sections show a distinct puncta (arrowhead) and Sgg protein signal in a bouton that hasn’t yet formed a postsynaptic density. (R) Bubble plot representing each NMJ with the percentage of GBs containing mRNA and protein on the X and Y axes respectively. Nearly 100% of GBs have Sgg protein with Area of the circle is proportional to the number of GBs per NMJ.

To address the degree to which mRNA localisation is likely to contribute to targeting proteins in the postsynapse, we characterised in detail the distribution of the 200 mRNAs and the proteins they encode at the muscle and postsynaptic cytoplasm. Our dataset shows that a large proportion of genes encode mRNAs that are present, but not enriched, within the postsynaptic density (PSD) without any corresponding protein enrichment or known synaptic function for the proteins encoded by the mRNAs (Supplementary Table S3). For example, the distribution of *nrv1* and *zap3* mRNAs are indistinguishable, even though the Nrv1 protein is strongly enriched at the PSD and Zap3 protein is evenly distributed throughout the muscle cell (Figure 7E-L). We identified 13 in total with strong enrichment at the PSD, none of which show obvious mRNA enrichment in the same location (Supplementary Table S3). Nevertheless, mRNA from a large proportion of genes is present at the PSD, despite not being enriched in that compartment relative to the rest of the muscle cytoplasm. We interpret these results as indicating that mRNA localisation does not play as strong a role in the NMJ postsynapse. Given that in the postsynaptic (muscle) compartment, translation factors, such as eIF4E, are known to regulate NMJ development and plasticity (Menon et al. 2004), we propose that spatial regulation of gene expression makes a strong contribution to the postsynapse, but through localised translation, as in the case of Msp300 (Titlow et al. 2020).

### Active *sgg* mRNA transport and activity-dependent accumulation of *sgg* in ghost boutons

Localisation of mRNA in the *Drosophila* NMJ axon terminal has not previously been demonstrated at the single molecule level. To determine whether *sgg* mRNA is actively regulated during plasticity at the larval NMJ, we performed a set of experiments using transport mutants and a spaced KCl stimulation paradigm (Ataman et al. 2008). We found that not only is *sgg* actively transported to the axon terminal, Sgg protein levels in the axon terminal are elevated in response to KCl stimulation, and both *sgg* mRNA and protein appear in newly formed synaptic boutons. We anticipate that at least a number of other transcripts we identified in the motoneuron axonal synapses will be similarly locally translated in response to neuronal activation. However, carrying out such experiments systematically for 69 genes (Supplementary Table S2) is beyond the scope of this study. Nevertheless, our demonstration of local translation of Sgg protein raises the possibility that many more low abundance mRNAs at the distal axonal synapses are locally translated.

Kinesin-1 is known to be required for transport in neurons in many circumstances and tissue types. To determine the mechanism for *sgg* mRNA localisation to axon terminals of the larval NMJ we carried out smFISH experiments on *kinesin1 heavy chain* mutant third instar larval NMJs. We found that *sgg* mRNA is actively transported to motor axon terminals by the kinesin-1 motor. The number of *sgg* mRNA molecules in axon terminals was measured by counting the number of diffraction limited fluorescent spots in the images of smFISH within a 3D-segmented axonal volume. *sgg* mRNA measurements were acquired in wild type larvae, and in a trans-heterozygous *khc* mutant (*khc*^*23*^ */ khc*^*27*^), a combination of an amorphic and a hypomorphic allele that avoids lethality. Loss of *khc* function resulted in an 84% reduction (Figure 8E) in the number of *sgg* transcripts per NMJ (Figure 8A-C), indicating that kinesin-1 based transport is required for *sgg* localisation at the NMJ.

We then asked whether kinesin-1 based transport is required for activity-dependent synaptic plasticity using a patterned chemical stimulation assay that induces the formation of new synaptic boutons, a form of activity-dependent synaptic plasticity that requires protein synthesis (Ataman et al. 2008). In this assay, boutons with presynaptic labelling that have not yet acquired the postsynaptic density marker Dlg1 are immature, so-called ‘ghost boutons’. Using KCl stimulation and ghost bouton labelling as a readout of structural plasticity, we found that loss of *khc* function resulted in an 85% reduction in the number of activity-induced ghost boutons (Figure 8F-H). Although loss of *khc* function also disrupts protein and organelle transport, these experiments show that *sgg* mRNA is actively transported to the synapse, and that kinesin-1 based transport is generally required for structural plasticity at the larval NMJ.

The canonical function of localised mRNA is to provide an immediate source of new protein translation in response to an external stimulus. To determine whether Sgg protein levels are elevated in response to patterned KCl stimulation, we quantified Sgg::YFP fluorescence levels in KCl-stimulated larval fillet preparations relative to mock-treated controls. We found a modest but highly significant increase in Sgg::YFP levels at stimulated NMJs (12.4 ± 0.02%; p = 0.0002, Student’s t-test; Figure 8I-K). It is perhaps not surprising to observe such a modest increase in protein level, given that the fluorescence intensity measurements were averaged across the entire axon terminal, while the response is expected to be highly localised. In fact, we often observe high signal intensity concentrated in individual boutons that are immature in appearance (Figure 8J, arrows), suggesting that Sgg protein is localised at higher concentrations in axon terminals during the early stages of synapse formation.

To determine if Sgg appears in newly formed synaptic boutons, we repeated the KCl stimulation experiment, and imaged fixed samples labelled with presynaptic and postsynaptic markers. Sgg protein was present in 93% of ghost boutons (out of 27 ghost boutons from 14 NMJs in 5 different animals; Figure 8L-R), indicating that Sgg is almost always present during the early stages of activity-dependent synapse formation. To ask whether localisation of *sgg* mRNA could play a role in the accumulation of Sgg protein in ghost boutons, these samples were also labelled with smFISH probes targeting *sgg mRNA*. We detected *sgg* mRNA in over 20% of ghost boutons (Figure 8L-R). We interpret our results as indicating that only some of the Sgg protein is translated from mRNA locally, whereas some Sgg protein is likely to be transported to the synapses in response to activation.

## Discussion

We present a data resource and a generalisable strategy to investigate the mechanisms of spatial gene expression control for a large number of gene candidates, at subcellular resolution, and across multiple whole tissues. To facilitate the extraction of new biological knowledge from this dataset, we have developed a computational pipeline to annotate and browse the image data, and systematically interrogate the imaging data alongside existing genomic and phenotypic datasets. This approach has yielded insight into post-transcriptional regulation that improves our understanding of both brain development and synaptic plasticity.

### A powerful method to quantify the entire gene expression lifecycle for any gene

Gene expression is a multistep process that is rarely investigated end-to-end, from transcription, to mRNA processing, to protein production. Our approach to measuring gene expression provides important insight into how an individual gene is regulated *in vivo*, while also highlighting the need to understand mechanisms of post-transcriptional regulation in more detail. This approach could be especially powerful in model organisms where large collections of protein traps are already available, and for tissue types in addition to the nervous system in *Drosophila*. We developed software tools that make it easier to systematically assemble, annotate and classify the imaging data for curation (Figure 2).

### Estimating the contributions of post-transcriptional regulation to brain development

Our dataset highlights a set of genes that exhibit obvious discordance between mRNA and protein expression levels throughout the neuroblast lineage in the larval central brain. This result is consistent with bulk sequencing studies that have identified large sets of genes that have mismatched levels of mRNA and protein (Y. Liu, Beyer, and Aebersold 2016; Buccitelli and Selbach 2020). We show, with high spatial resolution in a tissue-specific context, that lack of correlation between mRNA and protein levels often arises between cells at different stages of neuronal differentiation. Lack of correlation between mRNA and protein concentration in a cell could arise through many different mechanisms, which can be divided into two classes. First, mechanisms depending on the spatially distinct production of protein and second, mechanisms depending on differences in mRNA or protein decay. The quantitative power of our smFISH data can provide an estimate of mRNA synthesis and decay rates. Specifically, differences in the ratio of protein to mRNA provides an estimate of translation rates, and protein decay can be assumed to account for instances that are not explained by differences in mRNA metabolism or translation. By definition, each gene that we define as being post-transcriptionally regulated has equal levels of mRNA synthesis across the cell lineage. The occurrence of translational regulation highlights a major gap in our understanding of translational control, because it is not clear from our data whether these genes are more translationally active and/or translationally repressed across different stages of cell differentiation. Identifying the trans-acting factors present and their interactions with specific mRNAs at different stages of development, will be necessary to fully understand how these genes are regulated to influence neural differentiation. However, such studies can only be carried out for specific cases, and a global analysis of the trans-acting factors and their signals for the 200 genes we have characterised is beyond the scope of this study.

Our analysis revealed 12 out of 200 genes that showed obvious post-transcriptional regulation within the neuroblasts of the larval central brain (Figure 4 and Supplementary Table S3). An approximate extrapolation of that percentage to the whole genome indicates that over 500 genes are likely to exhibit similar expression patterns and post-transcriptional regulation in neuroblasts. The stability and translation of such mRNAs are known to be regulated by mRNA binding proteins. Syncrip and Imp are mRNA binding proteins that are already known to play a major role in *Drosophila* brain development (Samuels, Arava, et al. 2020; Samuels, Järvelin, et al. 2020; Z. Liu et al. 2015), with conserved mechanisms in the mammalian brain. However, Syp and Imp are certainly not unique, since out of the 523 known canonical mRNA binding proteins in *Drosophila (Sysoev et al*. *2016)*, 226 are expressed in neuroblasts (Berger et al. 2012). Characterising the function of so many mRNA binding proteins in neuroblast differentiation is daunting. Nevertheless, in the future high throughput approaches may have to be brought to bear on these large numbers of genes in order to understand the full complexity of the more than 500 genes estimated above to be likely to exhibit post-transcriptional regulation during brain development.

### The landscape of synaptic mRNA and protein localisation in an intact brain

Our study provides insight into the prevalence of synaptic mRNA localisation and local translation, which is thought to be a critical factor in synapse development (Shigeoka et al. 2016; Cioni, Koppers, and Holt 2018) and plasticity (Holt and Schuman 2013; Holt, Martin, and Schuman 2019). A series of elegant transcriptomic studies have revealed thousands of different mRNA species that are present in neurites—axons or dendrites—of various different neuronal cell types. A subset of those localised mRNAs make up a core set of neurite-enriched transcripts (von Kügelgen and Chekulaeva 2020), and localised mRNAs are likely to encode as much as half of the synaptic proteome in cultured neurons derived from mouse embryonic stem cells (Zappulo et al. 2017). We find that a slightly lower, but similar proportion of synaptic proteins are found alongside the mRNAs that encode them in the optic lobe, mushroom body, and sensorimotor neuropils of the *Drosophila* larval brain. On average, mRNAs detected in these *Drosophila* neuropils have mammalian orthologs that localise in at least 8 other synaptic transcriptome studies. Two of those genes, *ATPsynbeta* and *14-3-3epsilon*, are among the 10 most commonly detected mRNAs across the set of neurite transcriptome studies. The fact that these specific mRNAs are selectively localised to the neurite compartment, in different cell types and across millions of years of evolution, argues for the importance of their local translation in synaptic physiology.

Counterintuitively, mRNAs that encode nuclear proteins were highly enriched among the synaptic mRNAs in our dataset (Figure 5I-O). Retrograde signalling from synapse to nucleus is a relatively understudied process that contributes to many phases of the synaptic lifecycle, including development, plasticity, and response to injury (Cohen and Greenberg 2008; Fainzilber et al. 2011). Some of the signalling cascades that activate and execute synapto-nuclear signalling have been defined, as well as the transcription factors and genes that are upregulated in response to retrograde signalling. Consistent with synaptic transcriptome studies (von Kügelgen and Chekulaeva 2020), we detected several mRNAs at the synapse that encode transcription factors. These transcription factors could be translated in response to local changes in synaptic activity and trafficked back to the nucleus to induce the expression of long term memory genes. mRNAs encoding splicing factors Rm62 and qkr58E-1 were also detected at *Drosophila* synapses. Activity-dependent alternative splicing, like retrograde synapto-nuclear signalling, is poorly understood but known to be important for several phases of the synaptic lifecycle (Flavell et al. 2008; Hermey, Blüthgen, and Kuhl 2017). Our screen included some genes such as Rm62 and qkr58E-1, with functional connections to the spliceosome or splicing. The mammalian homologues of these two genes, Ddx17 and Khdrbs3 respectively, have mRNAs that are detected in at least 9 synaptic transcriptome studies (von Kügelgen and Chekulaeva 2020). Therefore, our data highlights the possibility that local synthesis and nuclear transport of Rm62 and qkr58E-1 could link elevated synaptic activity and the spliceosome.

### RNA localisation in glia and the neuromuscular junction

We provide the first evidence for mRNA localisation in the peripheral processes of *Drosophila* glia (Figure 6). Like neurons, glia have long cellular processes that exhibit mRNA localisation and local protein synthesis (Pilaz et al. 2016; Sakers et al. 2017), however the functional role of mRNA localisation in glia is not well defined. Our results show that several cell junction and membrane proteins are encoded by mRNAs that are localised in glial processes at the neuromuscular junction and in the central nervous system, cells which are functionally homologous to vertebrate Schwann cells and oligodendrocytes, respectively. Consistent with glial transcriptomic studies, which tend to show lower transcript diversity than neurite transcriptomes, we find that the relative number of genes expressed in glial processes is lower than in neural processes. We also find that the majority of proteins expressed in glial processes have localised mRNAs (88% across peripheral and nervous system glia, (Supplementary Table S3)), whereas the estimated contribution of local mRNA to the synaptic proteome in neurons is ∼50% (Zappulo et al. 2017). Together, these results suggest that mRNA localisation in glial processes is highly regulated, and is likely to make an important contribution to the local proteome. The extent to which glial processes influence synaptic transmission and activity-dependent plasticity at the larval neuromuscular junction is not yet known, but glial signalling through the Wnt pathway (Kerr et al. 2014) and Endostatin pathway (T. Wang et al. 2020) have been shown to disrupt synaptic physiology. Those signalling factors, in addition to the genes identified in our dataset, could be regulated by local translation at the neuromuscular synapse in response to extracellular cues. Moreover, the presence of localised mRNAs in peripheral processes of cortical and ensheathing glia suggest that such mechanisms could be important for cognitive function and brain development.

We were surprised by the absence of mRNA enrichment at the postsynaptic density (PSD) of the larval neuromuscular junction (Figure 7). Although 13 genes encode proteins that are highly enriched at the PSD (Supplementary Table S3), none display a corresponding enrichment of mRNA. Despite the lack of mRNA enrichment, we observed a high abundance of mRNAs at the PSD encoding many different types of proteins, an indication that local translation does occur. The data are consistent with a model where specificity of local translation, for example in response to elevated synaptic activity, is achieved through translational regulation by selective mRNA binding proteins, as shown previously for activity-dependent regulation of Msp300 by an hnRNP mRNA binding protein called Syncrip (Titlow et al. 2020). Here we report another conserved mRNA binding protein, RnpS1, which has the potential to provide highly localised regulation of mRNA dynamics, based on its strong enrichment in discrete punctate particles at the PSD (Figure 7S).

Our approach provides a framework for future studies aimed at understanding gene expression control, at scale, across 3D tissue landscapes with heterogeneous cell types. By surveying a small percentage of protein coding genes we undoubtedly underestimate rare expression patterns, but this diverse sample provides a useful estimate of the frequency in which the observed gene expression patterns are expected to occur genome-wide. Future studies can be expanded with the use of high throughput slide scanning microscopes and the extensive collection of protein trap lines available in *Drosophila (Nagarkar-Jaiswal et al. 2015)*. Additional cellular markers, for example axon and dendrite specific markers in synaptic neuropils, and additional developmental time points would also be valuable for identifying temporal changes in mRNA dynamics. This approach to investigating gene expression provides critical insight into how gene function is regulated within the tissue environment.

## Supporting information

Accompanying data for Zegami

## Acknowledgements

We are very grateful to the Bloomington, Vienna, and Kyoto Drosophila Stock Centres (fly stocks), Flybase and Flymine (Lyne et al. 2007) for their reagents and open data, which were invaluable to this work. We are grateful to David Ish-Horowicz, Alfredo Castello and members of the Davis laboratory for critical reading of the manuscript and feedback on the project. We thank Zegami Ltd. for their help, advice and hosting the collection. This work was generously supported by a Wellcome Senior Research Fellowship (096144) and Wellcome Investigator Award (209412) to I.D., which funded A.I.J. R.M.P., J.S.T., MK.T. Advanced microscopy facilities and technical advice as well as support to DM.SP. were provided by Micron Oxford (https://micronoxford.com), supported by Wellcome Strategic Awards (091911 and 107457) and a Medical Research Council/Engineering and Physical Sciences Research Council/Biotechnology and Biological Sciences Research Council next-generation imaging award to I.D. as the principal investigator. J.S.T. and MK.T. were supported by a Leverhulme Trust grant to I.D. Department of Biochemistry DPhil studentships supported J.Y.L. and D.S.G. M.K. was supported by the Biotechnology and Biosciences Research Council (BBSRC), grant numbers: (BB/M011224/1) and (BB/S507623/1), by A.G. Leventis Foundation and by Zegami Ltd.

## Author contributions

J.S.T. and I.D. conceived and designed the study, interpreted results, wrote and revised the manuscript as well as provided supervision and data quality control. J.S.T. performed experiments and generated many figures and analysed data. A.P. and J.Y.L. performed the majority of experiments, made many figures. M.K. generated figures, curated data, performed formal data analysis including conceiving the data annotation software, revised and edited the manuscript and created the Zegami instance. D.S.G. performed follow-up experiments. D.E., J.J.S.Y., F.L.Y, S.G., H.S.F., F.S., H.M., A.D-P., S.T., D.A. and I.K. performed experiments. D.M.S.P, M.K., MK.T. and J.Y.T. developed software for data annotation. D.E., J.Y.L., A.I.J. performed data analysis. S.T. helped with the Zegami instance creation and provided guidance. D.E. supervised experimental execution including data quality control and performed project management and coordination.

## Materials and Methods

### Animal model

*Drosophila melanogaster* stocks were maintained with standard cornmeal food at 25°C on 12-h light–dark cycles unless otherwise specified. Wandering third instar larvae were used for all experiments. The following genotypes were used: Canton S (wild type unless otherwise specified), YFP insertion lines were from the Cambridge Protein Trap Insertion project (Lowe et al., 2014), repo-Gal4 (Sepp, Schulte, and Auld 2001) and UAS-mcd8-mCherry (BDSC #27391)

### Whole-mount single molecule fluorescence in situ hybridization (smFISH) and immunofluorescence

*Drosophila* larval CNS and NMJ specimens were prepared using a protocol that was previously described (Titlow et al., 2018). Briefly, specimens were fixed in PFA (4% in PBS with 0.3% Triton X-100 [PBTX]) for 25 min, rinsed three times in PBTX, blocked for 30 min in PBTX + BSA (1%), and incubated overnight at 37°C in hybe solution (2× SSC, 10% formamide, 10% dextran-sulphate, smFISH probes [250 nm; individual probe sequences listed in Table S1], and primary antibodies). The next morning, samples were rinsed three times in smFISH wash buffer (2× SSC + 10% formamide) and incubated for 45 min at 37°C in smFISH wash buffer with secondary antibodies and DAPI (1 μg/ml), and then washed for 30 min in smFISH wash buffer at room temperature before mounting in glycerol (Vectashield). PBTX was used in place of smFISH wash buffer for experiments that did not require smFISH. The following antibodies were used: mouse anti-Dlg1 (1:500; 4F3, Developmental Studies Hybridoma Bank), HRP-Dyelight-405/488/Alexa Fluor 568/Alexa Fluor 659 (1:100; Jackson ImmunoResearch Laboratories), donkey anti-guinea pig Alexa Fluor 488 (1:500; Thermo Fisher Scientific), and donkey anti-mouse Alexa Fluor 568 (1:500; Thermo Fisher Scientific). We estimate that the efficiency of detection of individual mRNA molecules were 51% in the brain and 64% in the NMJ (Figure 1E’’). Although we anticipate that the exact efficiency of detection varies considerably between experiments, these efficiencies are likely to be conservative estimates, and in many individual experiments the sensitivity is likely to be higher. Measuring the sensitivity for every individual experiment is not practical.

### Image acquisition, post processing, and analysis

Whole-mount immunofluorescence and smFISH specimens were imaged on a spinning disk confocal microscope (Ultraview VoX; PerkinElmer) with 60× oil objective (1.35 NA, UPlan SApo, Olympus) and electron-multiplying charge-coupled device camera (ImagEM; Hamamatsu Photonics) or laser scanning confocal microscope (Olympus Fluoview 3000, 1.30 NA SI UPLASAPO60XS2, GaSP detector; or Zeiss LSM880, 63x 1.4 NA oil objective, GaSP detector). Background subtraction was accomplished with the rolling ball subtraction algorithm in ImageJ (radius = 5 pixels for smFISH data and radius = 20 pixels for cell markers).

### Spaced potassium stimulation protocol

Third instar *Drosophila* larvae were dissected in two separate chambers (35mm Sylgard elastomer-lined petri dishes) to allow even saline perfusion from peristaltic pumps. A series of five short high potassium saline (KCl, 90 mM) pulses (2, 2, 2, 4, and 6 min) were separated by 15-min perfusion of HL3 saline as described previously (Ataman et al., 2008)). For smFISH and immunofluorescence, the larvae were fixed 150 min after the first stimulus, and images were acquired on the spinning disk confocal system described in the section above. For live imaging experiments, images were acquired on the Zeiss LSM880 system described above (20x 1.0 dipping objective) from 10 min after the last stimulus.

### Software pipeline for browsing and annotating the dataset

We built a generalizable pipeline to display high resolution microscopy images, annotate specific features, and browse the collection from each gene together with relevant publicly available data. An overview of the pipeline is shown in Figure 2. Raw image data were uploaded to an OMERO server where multi-channel Figures displaying multiple fields of view (FOVs) were generated for each image and displayed in OMERO.Figure at the appropriate image plane. After creating a separate figure for each gene in various nervous system compartments, Figures were extracted as .jpg files that were used both for annotation and to build a browsable image analysis platform in Zegami. To enrich the image collection we added phenotypic and physical information corresponding to each gene, and expanded this information to include the whole genome in an effort to impute our screening results to genes with similar characteristics. Included in the pipeline is a Python script that extracts user-specified gene data from Intermine and local .csv files.

### Annotation comparison and conflict resolvement

Multiple members of the Davis lab with expert knowledge of the larval nervous system tissue and smFISH signal interpretation annotated the figures. Each figure was annotated by three different scorers and the annotations were compared using a Python script which we wrote in house. A majority vote was used to resolve any conflicts between the answers selected by the experts. When a majority view could not be reached, a fourth expert was required to resolve the specific conflict, by focusing with more time on the particular set of images alone. This approach ensured the high confidence and quality of all our annotations.

### Annotate.OMERO.Fig

To facilitate image annotation we built a Python application with a graphical user interface using the PyQt5 library. The app makes it easy to systematically cycle through a large set of images and score them based on a list of user-defined questions with True/False, multiple choice, or write-in answers. The output is a .csv file that can be directly analysed or uploaded together with phenotype and gene information to a Zegami database. The github repository submitted with this manuscript contains code that interfaces with the Intermine Python API to append flymine queries, and to local file directories to append various public datasets from different model organisms.

### Statistical analysis of GB, mRNA, and protein levels

Statistical tests that were applied to each dataset are given in the figure legends along with the number of samples appearing in each graph. The normality assumption was tested with the Shapiro–Wilk test. The equal variances assumption was tested with an F test or Levene’s test, depending on the number of groups. Normally distributed populations with equal variances were compared using Student’s t test or one-way ANOVA (with Tukey test for multiple comparisons), depending on the number of groups. Populations with nonnormal distributions were compared using the Wilcoxon rank sum test or Kruskal–Wallis test (with Dunn test for multiple comparisons), depending on the number of groups. All statistical analyses were performed in R (v3.3.2 running in Jupyter Notebook).

### Lead contact

Further information and requests for resources and reagents should be directed to and will be fulfilled by the lead contact, Ilan Davis.

### Data and Code Availability

All Python code is freely available at the GitHub repository: https://github.com/ilandavislab/Annotate.OMERO.Fig (copy archived at Zenodo here). A Zegami instance can be accessed here and the accompanying data are available in the Supplementary material. Guidance on how to use Zegami and browse the collection can be accessed here.

## FIGURES

**Figure S1.**
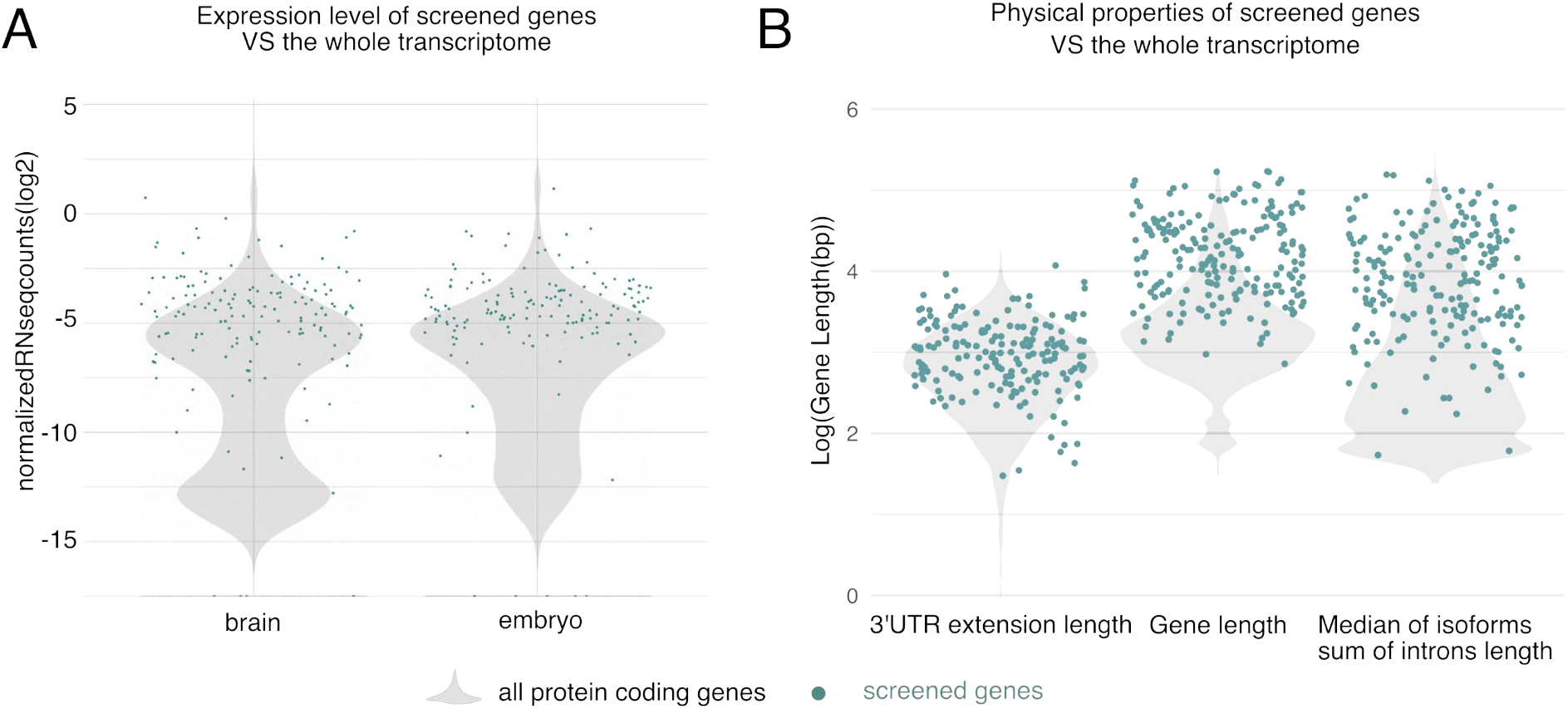
Expression and physical properties of screened genes are mostly representative of the whole transcriptome. Previously published datasets (Flymine) were used to determine if the collection of screened genes show any biases relative to the rest of the genome. (A) Distribution of expression levels in the screened genes are similar to the rest of the transcriptome in both the brain, and in the embryo. (B) While the length of 3’UTR extension in the screened genes is similar to the rest of the transcriptome, the screened genes on average have a longer total length and longer introns.

**Figure S6.**
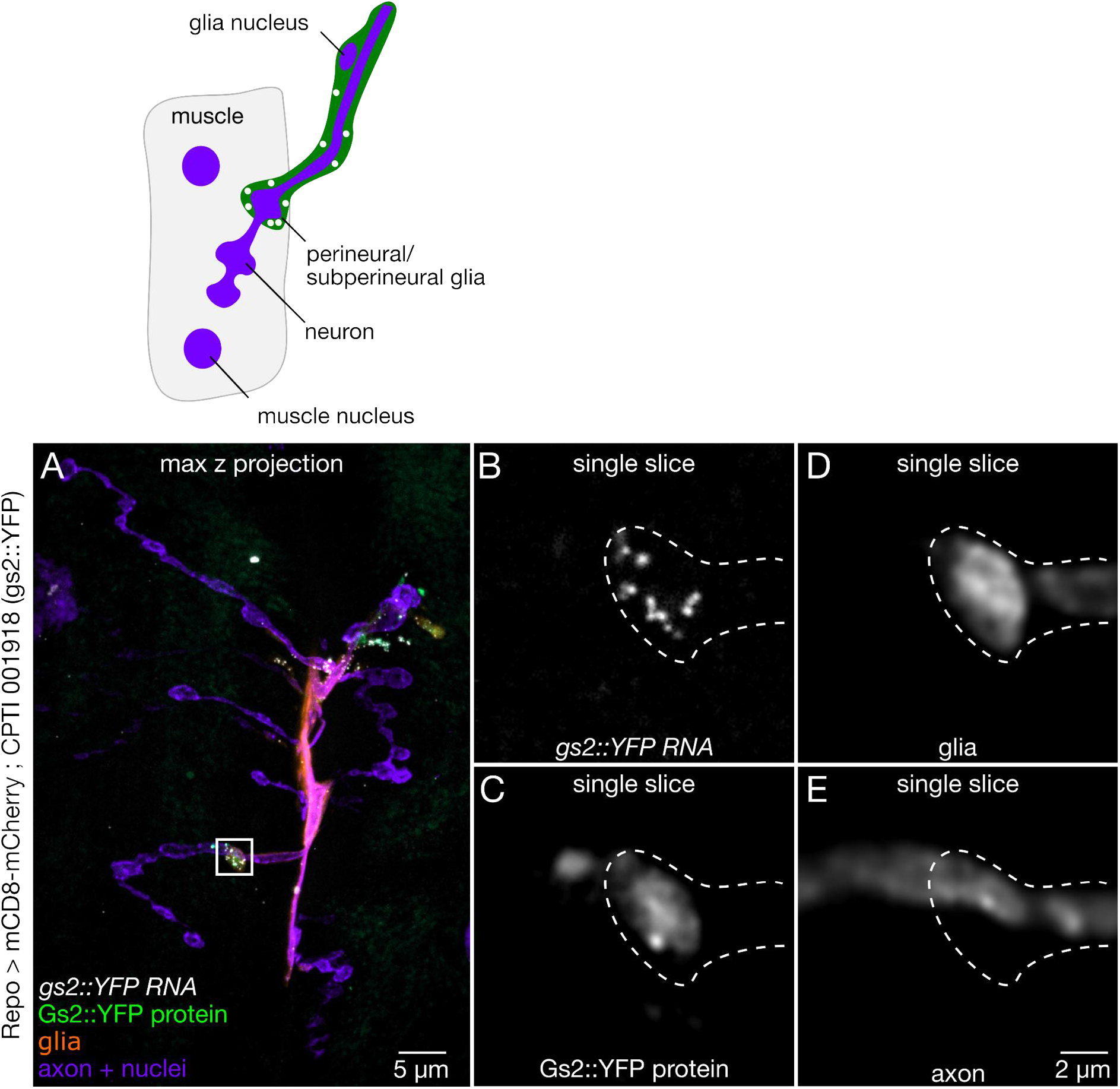
*Gs2 mRNA* is localised in glial boutons at the larval neuromuscular junction. Representative confocal images showing *gs2::YFP* protein and mRNA expression (white) at the NMJ with markers for glia (Repo > mCD8-mCherry, orange) and neurons (HRP, purple). (A) Full field of view image shows *gs2* expression is confined to proximal regions of the axon terminal where the glial reporter is located. (B-E) Magnified regions of a single optical slice from A (white box), showing several molecules of *gs2 mRNA* and protein in a perineurial glial process enveloping a bouton.

